# Evolution of CYP71D as a driving force of the diversification of monoterpene indole alkaloid biosynthesis

**DOI:** 10.1101/2025.01.09.632084

**Authors:** Sébastien Besseau, Mickael Durand, Duchesse-Lacours Zamar, Enzo Lezin, Louis-Valentin Méteignier, Caroline Birer Williams, Christelle Dutilleul, Thomas Perrot, Arnaud Lanoue, Audrey Oudin, Nicolas Papon, Chao Sun, Benoit St-Pierre, Vincent Courdavault

**Author notes:** Address correspondence to (S. Besseau) and (V. Courdavault). Present address: PHIM Plant Health Institute, Univ Montpellier, INRAE, CIRAD, Institut Agro, IRD, Montpellier, France.

## Abstract

Monoterpene indole alkaloids (MIAs) constitute a vast group of plant natural products synthesized within the Gentianales order. MIAs possess outstanding pharmacological properties that explain their wide use in the treatment of human diseases. These biological activities result from the complex structure of MIAs that originate from intricated biosynthetic pathways involving several enzyme families and notably cytochrome P450s. The early steps of MIA synthesis involve several P450s from the 71 clade, which catalyse the conversion of various reduced strictosidine aglycones into MIAs from the mavacurane, strychnane, akuammilane, sarpagane, and heteroyohimane groups. An extensive study of these P450 distribution in genomic resources reveals that they all belong to a single evolutionary lineage named GAS clade, restricted to the MIA producing Gentianales (Gelsemiaceae and Rauvolfioideae). A related but distinct P450 lineage (named sister GAS clade) was found in non-MIA producing Gentianales and species producing structurally distinct MIAs (Rubiaceae). Functional characterization of 36 members of the GAS clade revealed that these enzymes felt into four main groups of activity depending on substrate acceptance (ex: tetrahydroalstonine, geissoschizine, ajmalicine…) and the nature of the catalyzed reaction (aromatization, cyclisation). These characterizations also lead to the identification of a yohimbane aromatization activity for several of these P450s, increasing the number of MIA scaffolds associated with GAS activity to six. Lastly, the chronology of emergence of these GAS activities was assessed by using ancestral sequence reconstitution, establishing that the initial ancestor of the whole GAS clade exhibits a main alstonine synthase activity. Subsequent gene duplication events, combined with neofunctionalization, facilitated the progressive emergence of eight additional activities, variably distributed among the four GAS subgroups. This comprehensive enzyme characterization establishes the evolutionary trajectory of the GAS clade, demonstrating how its diversification has progressively shaped the chemical diversity of MIAs in Gentianales.

---

While primary metabolism is assumed to be highly evolutionarily conserved in plants, specialized metabolisms are more prone to active diversification to allow adaptation to changing environments. This notably results from core metabolic enzyme hijacking or gene duplication events that are highly prevalent in plants, as observed for cytochrome P450s. This class of enzymes indeed catalyzes a vast array of biochemical reactions and has been progressively recruited for the biosynthesis of defensive molecules such as monoterpene indole alkaloids (MIAs) in plants from the Gentianale order (St-Pierre et al., 2013). MIAs exhibit exceptional biological activities explaining their long-dated use by humans as anticancer of hypotensive drugs for instance. Along evolution, MIAs dramatically diversified with more than 3,000 different molecules belonging to distinct subfamilies including mavacuranes, strychnanes, akuammilanes, sarpaganes, yohimbanes and heteroyohimanes in the iconic model plants *Catharanthus roseus* or *Rauvolfia tetraphylla* (**Fig. 1A**) (O’connor and Maresh, 2006). All MIAs originate from the common precursor strictosidine and its downstream conversions involving (i) deglycosylation catalyzed by strictosidine glucosidase (SGD), (ii) reduction ensured by alcohol dehydrogenases and (iii) oxidation, oxygenation, aromatization or reconfiguration of the core MIA skeleton mediated by P450s from the CYP71D subfamily (Hamberger and Nak, 2013, Kulagina et al., 2022). P450s that use reduced strictosidine aglycones as substrates (e.g. geissoschizine; tetrahydroalstonine or ajmalicine) include geissoschizine cyclase (GC), geissoschizine oxidase (GO), sarpagane bridge enzyme (SBE), alstonine synthase (AS) and serpentine synthase (SS), and ensure the radiating diversification of MIAs (**Fig. 1A**). Interestingly, these P450s cluster in a monophyletic clade (**Supplemental Fig 1**), which was renamed hereafter as GAS, according to their first letter name (GO/GC; AS; SBE/SS). This GAS clade encompasses enzymes with distinct biochemical properties ranging from multiple substrate acceptance for specific product synthesis (AS/SS/SBE) to single substrate preference for the synthesis of multiple products (GO/GC) (Kulagina et al., 2022). Such a diversity thus suggests that a coordinated evolution has likely guided the emergence of the crucial GAS to direct MIA diversification, but knowledge about the underlying processes remains scarce.

**Figure 1:**
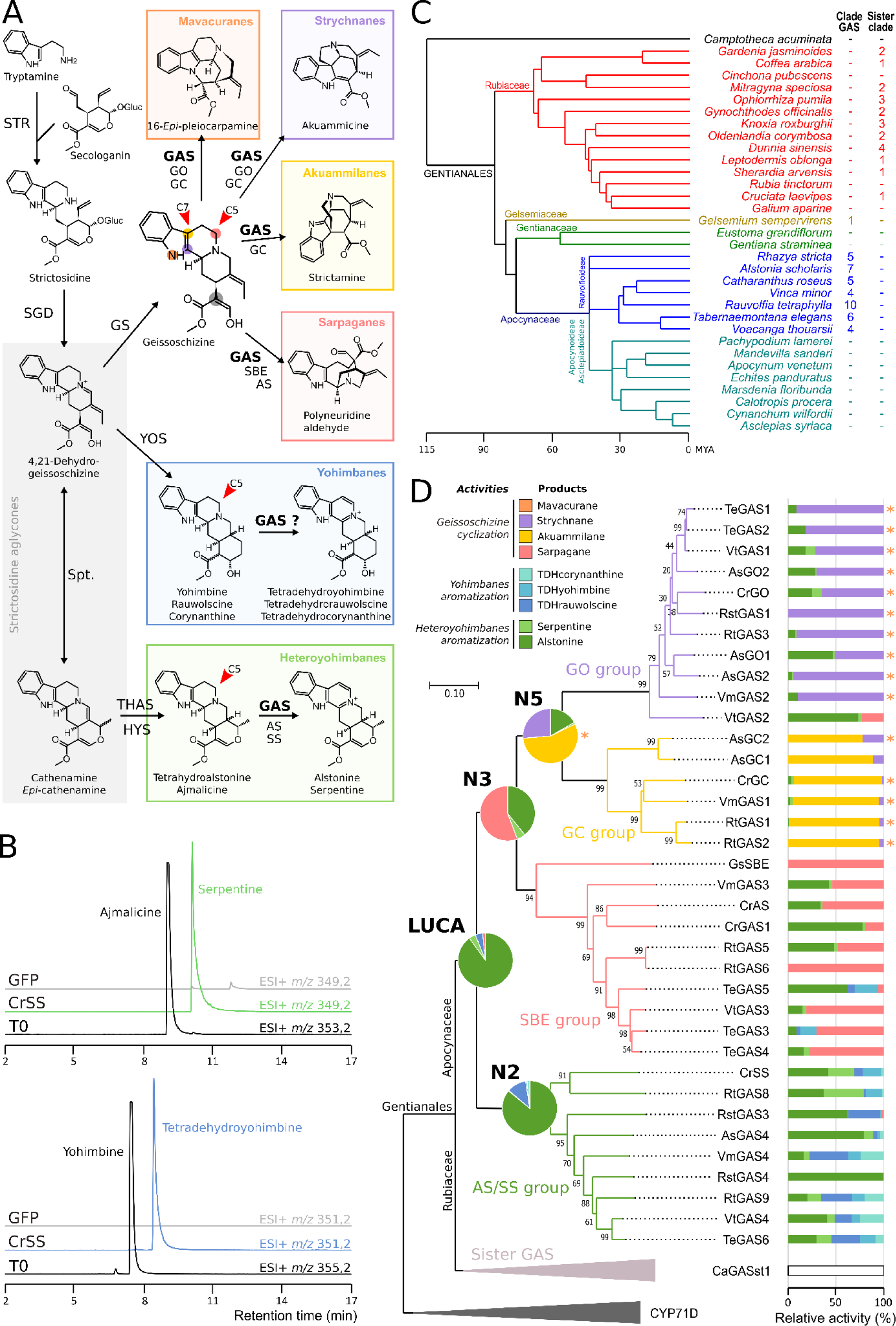
GAS distribution and evolution of their biosynthetic activity among Gentianales. **A.** Early steps of MIA biosynthesis pathways from strictosidine to MIA diversification (in colored boxes) conducted by GAS enzymes on the reduced strictosidine aglycones. Red arrows on molecules indicate the position of the initial oxygenation catalyzed by GAS, whereas the four colored dots on geissoschizine highlight positions of cyclization involving the keto-enol tautomer (gray dot) (see **Supplemental Fig 6** for details on the catalytic mechanism). STR, strictosidine synthase; SGD, strictosidine β-D-glucosidase; GS, geissoschizine synthase; YOS, yohimbane synthase; THAS, tetrahydroalstonine synthase; HYS, heteroyohimbane synthase; GO, geissoschizine oxidase; GC, geissoschizine cyclase; SBE, sarpagane bridge enzyme; AS, alstonine synthase; SS, serpentine synthase. **B.** CrSS was transiently expressed in *N. benthamiana* leaves that were further incubated with yohimbine (bottom panel). The reaction was analyzed by UPLC-MS in positive mode. The substrate was observed before incubation (T0) on ion chromatograms at m/z 355.2 (black trace) and the reaction product was monitored at *m/z* 351.2 (blue chromatogram) for the expected tetradehydroyohimbine. Leaf expressing GFP was used as a control in gray. The top panel presents the control activity of CrSS converting ajmalicine (*m/z* 353.2) into serpentine (*m/z* 349.2). **C.** Phylogeny of the Gentianales species used to study GAS distribution in plants. **D.** GAS members cloned and tested for activities were categorized using a neighbor-joining phylogenetic tree with 1000 bootstrap replications (%). Tobacco leaf disks expressing GAS candidates were separately incubated in a buffer containing 55 µM of tetrahydroalstonine or ajmalicine to evaluate heteroyohimbane aromatization, rauwolscine, yohimbine or corynanthine to evaluate yohimbane aromatization, and geissoschizine to evaluate its cyclization. After 24 hours, leaf disks were removed and products released in the incubation buffer were analyzed by LC-MS (typical activities are shown in **Supplemental Fig 3**). Histograms represent the GAS relative activities in percentage of cumulated product measured for each enzyme and activities of reconstructed ancestral sequences were shown on respective nodes. Traces of mavacurane synthesis are mentioned with an asterisk. GAS sub-clades are colored according to the main product obtained. The sister GAS clade and the CYP71D used to root the tree are condensed in light and dark gray respectively.

To gain insights into this evolutionary path, we first searched whether members of the GAS would be able to perform the aromatization of yohimbanes since this activity has never been reported before, while the corresponding compounds could be detected *in planta* (**Supplemental. Fig 2A-B**). We selected SS from *C. roseus* (CrSS) as a potential candidate since this enzyme performs a similar reaction on the related heteroyohimbane ajmalicine (Yamamoto et al., 2021). The corresponding gene was transiently expressed in *Nicotiana benthamiana* and the transformed leaves were fed with ajmalicine or yohimbine for assaying enzyme activity as compared to leaves expressing the green fluorescent protein (GFP). Analysis of the reaction products showed the expected aromatization of ajmalicine into serpentine while no modification was observed with GFP (**Fig. 1B**). More interestingly, we also observed the conversion of yohimbine into a putative tetradehydroyohimbine thus suggesting that CrSS also catalyzes aromatization of yohimbanes.The same reaction was observed with the yohimbine isomers rauwolscine and corynanthine (**Supplement Fig 3**).

We next searched for the whole set of GAS by analyzing the high-quality genomic sequences available in Gentianales (**Fig. 1C, Supplemental Table 1**). Interestingly, important differences were observed in the GAS distribution between the Gentianales families and subfamilies. Among Apocynaceae, GASs were systematically found in Rauvolfioideae, which is the main MIA producer subfamily, together with a large expansion of their members (4 up to 10). Conversely, no GAS homologs were found in Apocynoideae and Asclepiadaceae, thus suggesting that GAS-encoding genes were not retained during the evolution of Apocynaceae subfamilies that do not synthesize MIAs. Besides Apocynaceae, a unique GAS member was identified in Gelsemiaceae whereas only a GAS-related outgroup clade, hereafter named sister GAS, expanded among Rubiaceae that gathers non-MIA producing species and species producing structurally distinct MIAs such as quinoline alkaloids (camptothecin or cinchonine) (**Supplemental Fig 4**). Lastly, GAS and sister GAS were never jointly present in the same species and were absent outside of Gentianales, suggesting that both clades evolved from a common ancestral gene around 85 million years ago (**Fig. 1C**). This resulted in the acquisition of divergent functions including, among Apocynaceae, Gelsemiaceae and probably Loganiaceae, the specification for MIA biosynthesis we next further explored.

To evaluate the conservation of GAS member activities among Apocynaceae, 36 GASs out of the 45 identified from genomic resources were successfully cloned, besides the sister GAS sequence identified in the Rubiaceae *Coffea arabica* (CaGASst1). Each gene was transiently expressed in *N. benthamiana* leaves that were next fed with tetrahydroalstonine, ajmalicine, yohimbine, rauwolscine, corynanthine or geissoschizine as GAS reported substrates. The synthesis of the typical GAS products was monitored by LC-MS including heteroyohimbane aromatization (alstonine, serpentine), the newly identified yohimbane aromatization (tetradehydroyohimbine, tetradehydrorauwolscine or tetradehydrocorynanthine) and lastly geissoschizine cyclization leading to the synthesis of polyneuridine aldehyde (sarpagane scaffold), strictamine (akuammilane scaffold), akuammicine (strychnane scaffold) or 16-*epi*-pleiocarpamine (mavacurane scaffold) (**Fig. 1D**, **Supplemental Fig 3**). Firstly, while no activity was detected for CaGASst1 (sister GAS), we observed that all tested GASs display MIA biosynthesis activities that split into four main profiles according to the four enzyme subclades. The first one (AS/SS group), including 9 members such as CrSS and CrAS, catalyzes yohimbane and heteroyohimbane aromatization at different levels but has no activity on geissoschizine (Dang et al., 2018, Yamamoto et al., 2021). The second subclade (SBE group) with 10 members, including GsSBE and CrAS, mainly produces sarpagane derivatives but also alstonine to a lesser extent and serpentine traces (Dang et al., 2018). The third subclade (GC group), including AsGC1 and 2 among 4 other members, mainly converts geissoschizine into akuammilane and strychnane to a lesser extent, but also synthesizes traces of mavacurane (Méteignier et al., 2024). The last sub-clade (GO group) including CrGO mainly produces strychnane, various amounts of alstonine/serpentine and traces of mavacurane (Tatsis et al., 2017; Kamileen et al., 2024).

To figure out the evolution and the chronology of the emergence of these activities among the four GAS sub-clades, we reconstructed the most likely ancestral sequence for each node of the phylogenetic tree (**Supplemental Fig 4**). The oldest node belonging to the last universal common ancestor of GAS (LUCA), together with the ancestor of AS/SS group (N2), and the two ancestors from which diverged the SBE, GC and GO groups (N3 and N5) were thus selected for resurrection and activity assays using the whole set of precursors. Our results strongly suggest that the first GAS (LUCA) displays an original activity of alstonine synthesis by aromatization of tetrahydroalstonine that has probably been the first reduced strictosidine aglycone to emerge, in agreement with the multiplication of the genes encoding tetrahydroalstonine synthases (Stavrinides et al., 2016). Following an initial duplication event, one gene copy (N2) leading to the SS/AS group conserved this activity and slightly evolved to accept the ajmalicine isomer and yohimbanes as substrates (**Fig. 1D**). The substrate specificities of the AS/SS group members further evolved subsequently depending on species. The second copy (N3) gave birth to enzymes with a broader substrate acceptance, capable of cyclizing geissoschizine to synthesize sarpagane (SBE activity) while maintaining the initial AS activity. Interestingly, along with the acquisition of this new substrate acceptance, the enzyme catalytic mechanism remains conserved and still processing through C5 oxygenation of the substrate. This results in geissoschizine cyclization rather than aromatization since it contains a strong internal nucleophile keto-enol tautomer side chain at C16 thus allowing the formation of the new scaffold (**Fig. 1A**, **Supplemental Fig 5**). Later, a second duplication event led to the appearance of a gene encoding enzymes retaining the previous properties (SBE group) but also a second copy more prone to evolution (N5). The resulting enzymes (the GO/GC ancestor) still accepted tetrahydroalstonine as substrate but increased preference for geissoschizine and shifted its cyclization from sarpagane to strychane, akuammilane and mavacurane, as a possible consequence of the relocation of the oxygenated carbone on C7 (Méteignier et al., 2024). These activities were retained in the two last groups arising from a final local gene duplication event (Méteignier et al., 2024), resulting in the GC and GO groups, which slightly gained specialization for akuammilane and strychnane synthesis, respectively.

In conclusion, beyond characterizing the yohimbane aromatization activity, we have established that the evolutionary trajectory of the GAS has progressively shaped the diversity of MIA synthesized in Gentianales. Following the apparition of the initial AS activity, successive gene duplication events combined with neofunctionalization have progressively resulted in a broader substrate acceptance and transition of the GAS catalytic mechanism from tetrahydroalstonine aromatization to single and multiple geissoschizine cyclizations for a radiating diversification of MIAs. This originated from noticeable mutations of key residues in the GAS catalytic sites (**Supplemental Figure 6**) whose hierarchized substitutions would pave the way for future enzyme engineering.

## Accession numbers

CrGC (PP908688), CrGO (MF770508), CrAS (AHK60849), CrSS (MT829151), AsGC1 (OM323329), AsGC2 (OR604296), AsGO1 (OM323328), AsGO2 (PP908689), GsSBE (MF537712), CrGAS1 (PQ775227), VmGAS1 (PQ775228), VmGAS2 (PQ775229), VmGAS3 (PQ775230), VmGAS4 (PQ775231), RtGAS1 (PQ775232), RtGAS2 (PQ775233), RtGAS3 (PQ775234), RtGAS5 (PQ775235), RtGAS6 (PQ775236), RtGAS8 (PQ775237), RtGAS9 (PQ775238), AsGAS2 PQ775239(), AsGAS4 (PQ775240), VtGAS1 (PQ775241), VtGAS2 (PQ775242), VtGAS3 (PQ775243), VtGAS4 (PQ775244), RstGAS1 (PQ775245), RstGAS3 (PQ775246), RstGAS4 (PQ775247), TeGAS1 ( PQ775248), TeGAS2 (PQ775249), TeGAS3 (PQ775250), TeGAS4 (PQ775251), TeGAS5 (PQ775252), TeGAS6 (PQ775253), CaGASst1 (PQ775254).

## Author contributions

S.B., B.St-P and V.C. conceived and supervised the project. M.D. L.-V.M. and C.S. performed bioinformatics analysis. C.B.W. and A.L. performed metabolic analysis. S.B., D.-L.Z., E.L., C.D., T.P, A.O. and N.P. performed cloning, plant transformation and enzymes bioassays. S.B. and V.C. wrote the draft manuscript. All authors have reviewed and edited the manuscript.

## Acknowledgment

We would like to thank the chemical ecology platform of the Institut de Recherche sur la Biologie de l’Insecte, UMR 7261 CNRS, for the UPLC-Q-TOF services and facilities.

## Supplemental Information

### Online Methods

#### Genome mining

GAS-like sequences were retrieved from the publicly available genomes of 32 Gentianales (**Supplemental table 1**) through tBLASTn analysis (BLAST+ software, Camacho et al., 2009) using AsGC2, CrGO, GsSBE and CrSS as query. A threshold of 1e-10 E-value and 20% query coverage was applied. The translated result sequences were aligned with known GAS (AsGC, AsGC2, AsGO1, AsGO2, CrGC, CrGO, GsSBE, CrAS and CrSS) and a random list of 150 cytochrome P450s of CYP71D family, using ClustalW (Mega 11 software, Tamura et al., 2021). The phylogenetic tree was inferred using neighbor-joining to select sequences of interest depending on their placement close to GAS within the phylogeny. Genomic regions of candidates were explored to retrieve a complete coding sequence using tBLASTn analysis with the closest GAS homologue. Phylogeny of plant species was performed according to Time Tree of Life web server (Kumar et al., 2017).

#### GAS cloning

The full-length sequences of the candidate genes were amplified from cDNA or alternatively from gDNA using the Phusion High-Fidelity DNA polymerase and specific primers introducing BsaI or BsmBI restriction sites (**Supplemental table 2**). PCR products were purified with Nucleospin PCR clean-up kit and cloned into the pHREAC vector or a pHREAC version containing BsmBI sites (Lezin et al., 2024), using BsaI-HF-v2 or BsmBI-v2 golden gate assembly kits. The golden gate reactions were carried out in 10µl containing the T4 DNA ligase buffer 1X, 50 ng of vector, 1:3 molar ratio of purified PCR products and 10 units of golden gate mix. Reaction mixtures were incubated 10 cycles of 2 min at 37°C and 5 min at 16°C 10, followed by 5 min at 60°C and 5 min at 80°C. *E. coli* TOP10 was transformed with the Golden gate reaction products and selected on LB agar plates with kanamycin (50 µg/mL). Plasmids were extracted using Nucleospin plasmid kit and positive clones were sequenced.

#### Bioconversion assays

Electrocompetent *Agrobacterium tumefaciens* strain GV3101 cells were transformed with pHREAC vectors and selected on LB-agar plates supplemented with kanamycin (50 µg/mL) and rifampicin (50 µg/mL). Liquid cultures (10mL) supplemented with antibiotics were inoculated at 1/100 with pre-culture and incubated overnight at 28°C. Cells were harvested by centrifugation at 3000 g, resuspended in infiltration buffer (MES 10 mM pH 5.6, MgCl2 10 mM, acetosyringone 100 µM) to reach an OD600 of 0.2, and incubated 3h at 28°C under agitation before infiltration into younger leaves of 4 week-old *N. benthamiana* using syringes. Four days after infiltration, groups of 5 leaf disks of 1 cm diameter were collected and distributed into 24-well plates containing 750 µl of infiltration buffer with 55 µM substrate (tetrahydroalstonine, ajmalicine, rauwolscine, yohimbine, corynanthine or geissoschizine). After infiltration under vacuum at 50 mBar, plates were incubated in a greenhouse. After 24h, the medium was collected and diluted with 4 volumes of methanol prior LC-MS analysis. For akuammilane detection, the medium was alkalinized to pH 11 with NaOH 1 M before dilution in methanol.

#### LC-MS analysis

Reaction products were analyzed on a UPLC system (Acquity, Waters) coupled to a single quadrupole mass spectrometer (SDD2, Waters) equipped with an electrospray ionization (ESI) source, operating in positive ion mode. Samples were separated on a Waters Acquity HSST3 C18 column (150 x 2.1, 1.8 µm) with a flow rate of 0.4 mL/min at 55°C, using a 18-min linear gradient from 10 to 40% acetonitrile containing 0.1% formic acid. The capillary and sample cone voltages were 3000 V and 30 V, respectively. MS detection was carried out in the selected ion monitoring mode targeting [M + H]+ of products attempted. Geissoschizine (*m/z* 351.2, Rt 8.9 min), strictamine (*m/z* 323.2, Rt 8.0 min), strictamine isomer (*m/z* 323.2, Rt 6.8 min), akuammicine (*m/z* 323.2 Rt 9.0 min), polyneuridine aldehyde (*m/z* 353.2, Rt 9.5 min), 16-*epi*-pleiocarpamine (*m/z* 323.2, Rt 10.1 min), tetrahydroalstonine (*m/z* 353.2, Rt 9.4 min), ajmalicine (*m/z* 353.2, Rt 9.3 min), alstonine (*m/z* 349.2, Rt 10.5 min), serpentine (*m/z* 349.2, Rt 10.3 min), rauwolscine (*m/z* 355.2, Rt 6.2 min), yohimbine (*m/z* 355.2, Rt 7.7 min), corynanthine (*m/z* 355.2, Rt 7.9 min), tetradehydrorauwolcine (*m/z* 351.2, Rt 7.6 min), tetradehydroyohimbine (*m/z* 351.2, Rt 8.6 min), tetradehydrocorynanthine (*m/z* 351.2, Rt 9.7 min). MS2 acquisitions were performed on UPLC system (Acquity, Waters) coupled to a quadripole time-of-flight mass spectrometer (Synapt G2-Si HDMS, Waters) equipped with an electrospray ionization (ESI) source, operating in positive ion mode. Samples were separated on a Waters Acquity BEH C18 column (100 x 2.1, 1,7 µm) with a flow rate of 0.4 mL/min at 40°C, using a 12-min linear gradient from 10 to 60% acetonitrile containing 0.1% formic acid. Samples were analyzed in fast data-dependent acquisition (fDDA) mode consisting of a full MS survey scan from 50 to 1200 m/z (scan time = 0.1 msec) followed by MS/MS scans for the three most intense ions (m/z 100`–`1200; scan time = 0.1 msec). A collision energy ramp was set from 10 to 40 eV for low masses and 40 to 90 eV for high masses. Comparison of MS2 spectra were processed with R version 4.4.1 with the package Spectra (Rainer et al., 2022). Spectra were filtered by removing all peaks with an intensity smaller than 1% of the maximum intensity of each spectrum, and then peaks of the two spectra were matched with the same approach used in GNPS.

#### Ancestral sequence reconstruction

ASR analyses were performed using protein sequences of the 67 GAS-like identified among Gentianales genomes (GAS and sister GAS outgroup). The cytochrome P450 N-terminal transmembrane domain was removed until the conserved motif PPGP. Multiple sequence alignments were then performed with ClustalW (Mega 11 software, Tamura et al., 2021) and a preliminary neighbor-joining tree was inferred to establish relationships between sequences leading to the identification of 5 distinct sub-clades. GBLOCK (Castresana, 2000) was used to identify and resolve indels not conserved between related sequences. Only five short insertion sequences specific and strictly conserved within sub-clades were identified and kept (**Supplemental figure 6**). A phylogenetic tree was inferred from MSA of curated sequences using maximum likelihood method with 1000 bootstrap replicates under Mega 11 software (**Supplemental figure 4**). This tree was rooted with the introduction of several CYP71D involved in MIA synthesis (T3O, T16H, TEX, PYS, T19H and VH). The tree and MSA were finally used for ASR with FastML using marginal reconstruction (Ashkenazy et al., 2012). The most likely ancestral sequence for each node of interest was reconstructed based on the highest probability at each residue site (**Supplemental table 3**) whereas the previously identified sub-clade specific insertions were removed depending of nodes. Finally, the T16H2 N-terminal transmembrane domain (Besseau et al., 2013) was reintroduced in the sequences. To resurrect ancestral proteins, codon-optimized nucleic acid sequences were synthetized, cloned into pHREAC vector, and expressed in *N. benthamiana*.

## Supplemental Figures

**Supplemental figure 1:**
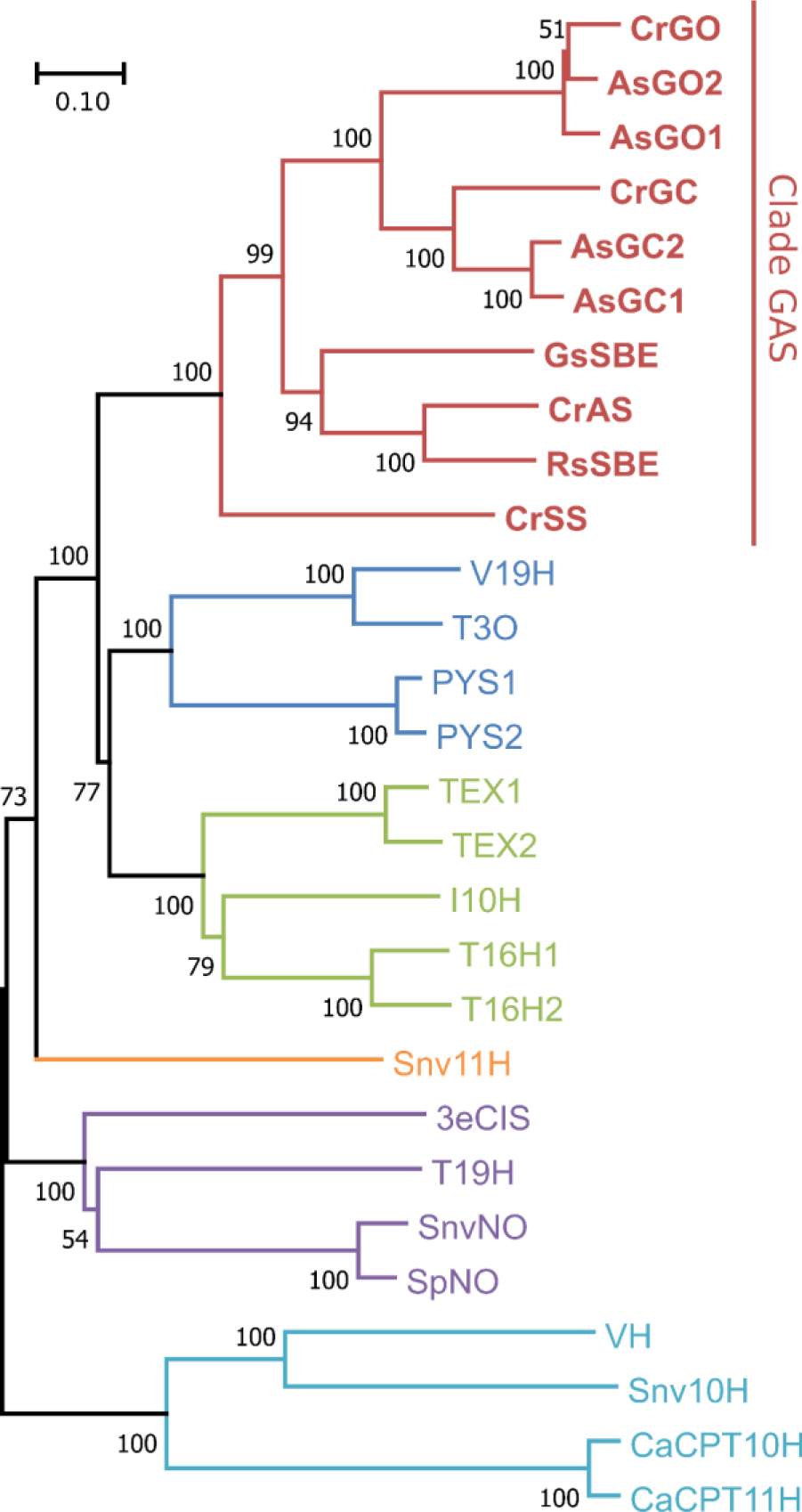
Phylogenetic analysis of cytochrome P450s involved in MIA biosynthesis. Protein sequences were categorized using a neighbor-joining tree with 100 bootstrap replications (the scale bar represents 0.1 substitutions per site). Enzymes belong to the CYP71D sub-family and form six monophyletic clades highlighted in colors. The clade renamed as GAS in this study appears in red. GO, geissoschizine oxidase; GC, geissoschizine cyclase; SBE, sarpagane bridge enzyme; AS, alstonine synthase; SS, serpentine synthase; V19H, vincadifformine 19 hydroxylase; T3O, tabersonine 3-oxygenase; PYS, pachysiphine synthase; TEX, tabersonine epoxidase; I10H, ibogaine 10-hydroxylase; T16H, tabersonine 16-hydroxylase; 11H, colubrine 11-hydroxylase; 3eCIS, 3-epi-corynoxeine/isocorynoxeine synthase; T19H, tabersonine 19-hydroxylase; NO, norfluorocurarine oxidase; VH, vinorine hydroxylase; 10H, strychnine 10-hydroxylase; CPT10H, camptothecin 10-hydroxylase; CPT11H, camptothecin 11-hydroxylase; Cr, *Catharanthus roseus*; As, *Alstonia scholaris*; Gs, *Gelsemium sempervirens*; Rs, *Rauvolfia serpentina*; Snv, *Strychnos nux-vomica*; Sp, *Strychnos potatorum*; Ca, *Camptotheca acuminata*.

**Supplemental figure 2:**
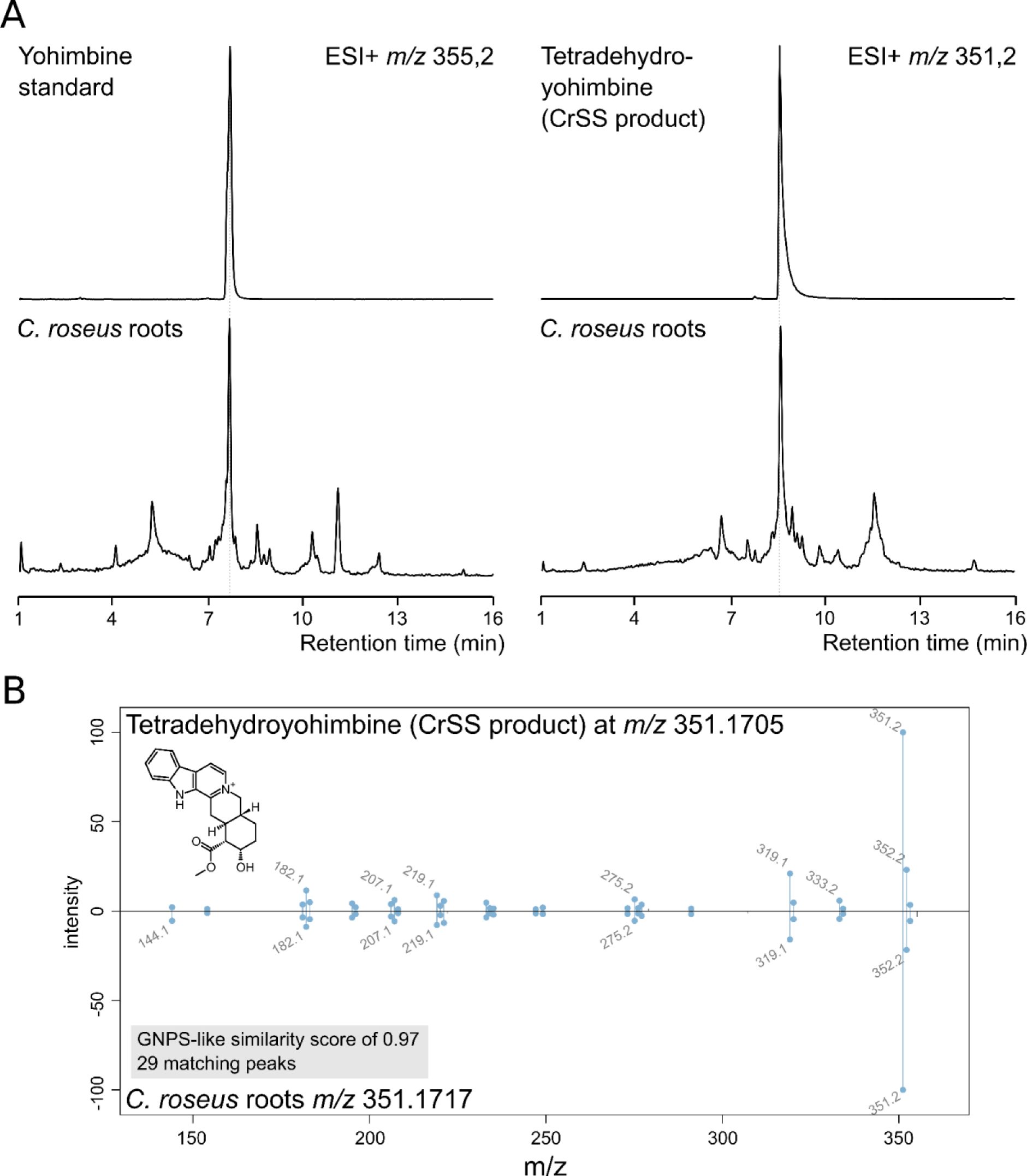
Yohimbanes and aromatized yohimbanes in *Catharanthus roseus* roots. A. Biological samples were analyzed by LC-MS and ion chromatograms in positive mode at *m/z* 355,2 and 351,2 were compared to authentic yohimbine standard and CrSS product assigned as 3, 4, 5, 6-tetradehydroyohimbine. B. Co-eluted compounds at *m/z* 351,2 were subjected to MS2 analysis on qTOF to confirm their similarity.

**Supplemental table 1:** List of genomic resources used to search for GAS.

| BioProject |  | Assembly |  |  | Organism Name | Assembly release Date |
| --- | --- | --- | --- | --- | --- | --- |
| Database | ID | Database | Accession | Name |  |  |
| NCBI | PRJNA476646 | NCBI | GCA_019593545.1 | ASM1959354v1 | Apocynum venetum | 13/08/2021 |
| NCBI | PRJNA787127 | NCBI | GCA_002018285.1 | ASM201828v1 | Asclepias syriaca | 03/03/2017 |
| NCBI | PRJNA292713 | NCBI | GCA_004801955.1 | ASM480195v1 | Calotropis procera | 22/04/2019 |
| NCBI | PRJNA907167 | NCBI | GCA_028651295.1 | ASM2865129v1 | Catharanthus roseus | 14/02/2023 |
| NCBI | PRJNA865567 | NCBI | GCA_025175665.1 | Cinchona pubescens_v1.0 | Cinchona pubescens | 13/09/2022 |
| NCBI | PRJNA497895 | NCBI | GCA_003713225.1 | Cara_1.0 | Coffea arabica | 08/11/2018 |
| NCBI | PRJEB70271 | NCBI | GCA_963678965.1 | daCruLaev1.1 | Cruciata laevipes | 06/12/2023 |
| NCBI | PRJNA742435 | NCBI | GCA_025388415.1 | GRF_CWil_1.0 | Cynanchum wilfordii | 23/09/2022 |
| NCBI | PRJNA649408 | NCBI | GCA_041381385.1 | Dunniia sinensis_1.0 | Dunniia sinensis | 21/08/2024 |
| NCBI | PRJNA1026900 | NCBI | GCA_035221335.1 | ASM3522133v1 | Echites panduratus | 08/01/2024 |
| NCBI | PRJDB12119 | NCBI | GCA_036186865.1 | ASM3618686v1 | Eustoma grandiflorum | 02/09/2023 |
| NCBI | PRJEB76009 | NCBI | GCA_964106835.1 | daGalApar1.hap1.1 | Galium aparine | 06/06/2024 |
| NCBI | PRJNA477438 | NCBI | GCA_013103745.1 | ASM1310374v1 | Gardenia jasminoides | 15/05/2020 |
| NCBI | PRJNA1020452 | NCBI | GCA_036785695.1 | ASM3678569v1 | Gentiana straminea | 22/02/2024 |
| NCBI | PRJNA625354 | NCBI | GCA_020080225.1 | ASM2008022v1 | Gynochthodes officinalis | 24/09/2021 |
| NCBI | PRJNA985238 | NCBI | GCA_030378225.1 | ASM3037822v1 | Knoxia roxburghii | 30/06/2023 |
| NCBI | PRJNA552650 | NCBI | GCA_016801395.1 | O1 | Leptodermis oblonga | 04/02/2021 |
| NCBI | PRJNA802340 | NCBI | GCA_024706555.1 | ASM2470655v1 | Mandevilla sanderi x Mandevilla atrovioleacea | 17/08/2022 |
| NCBI | PRJNA933576 | NCBI | GCA_035583425.1 | ASM3558342v1 | Marsdenia floribunda | 11/01/2024 |
| NCBI | PRJNA849364 | NCBI | GCA_024721245.1 | ASM2472124v1 | Mitragyna speciosa | 22/08/2022 |
| NCBI | PRJEB50026 | NCBI | GCA_949775105.1 | Ocorymbosa-M15.v1 | Oldenlandia corymbosa var. corymbosa | 07/04/2023 |
| NCBI | PRJDB8685 | NCBI | GCA_016586305.1 | Opu_r1.4 | Ophiorrhiza pumila | 09/12/2020 |
| NCBI | PRJNA997810 | NCBI | GCA_040940245.1 | UniT_Pla_v1.0 | Pachypodium lamerei | 29/07/2024 |
| NCBI | PRJNA771251 | NCBI | GCA_030512225.1 | ASM3051222v1 | Rauvolfia tetraphylla | 20/07/2023 |
| NCBI | PRJNA291945 | NCBI | GCA_001752375.1 | RHA1.0 | Rhazya stricta | 04/10/2016 |
| NCBI | PRJNA976901 | NCBI | GCA_032355605.1 | ASM3235560v1 | Rubia tinctorum | 04/10/2023 |
| NCBI | PRJEB59033 | NCBI | GCA_948330725.1 | daSheArve1.1 | Sherardia arvensis | 26/01/2023 |
| NCBI | PRJNA873287 | NCBI | GCA_026122235.1 | UniT_Vmin_1.1 | Vinca minor | 10/11/2022 |
| NCBI | PRJNA860765 | NCBI | GCA_026120185.1 | UniT_Voth_1.1 | Voacanga thouarsii | 10/11/2022 |
| NCBI | PRJNA1124898 | NCBI | Under release | Under release | Tabernaemontana elegans | Under release |
| NCBI | PRJNA361128 | Dryad | n/a | cac_genome_assembly_v2.4 | Camptotheca acuminata | 18/07/2018 |
|  |  | Dryad | n/a | gel_v1_asm | Gelsemium semperviens | 25/10/2018 |
| CNGBdb | CNP0002381 | CNGBdb | CNA0069587 | A.scholaris.contig | Alstonia scholaris | 23/02/2023 |

**Supplemental figure 3:**
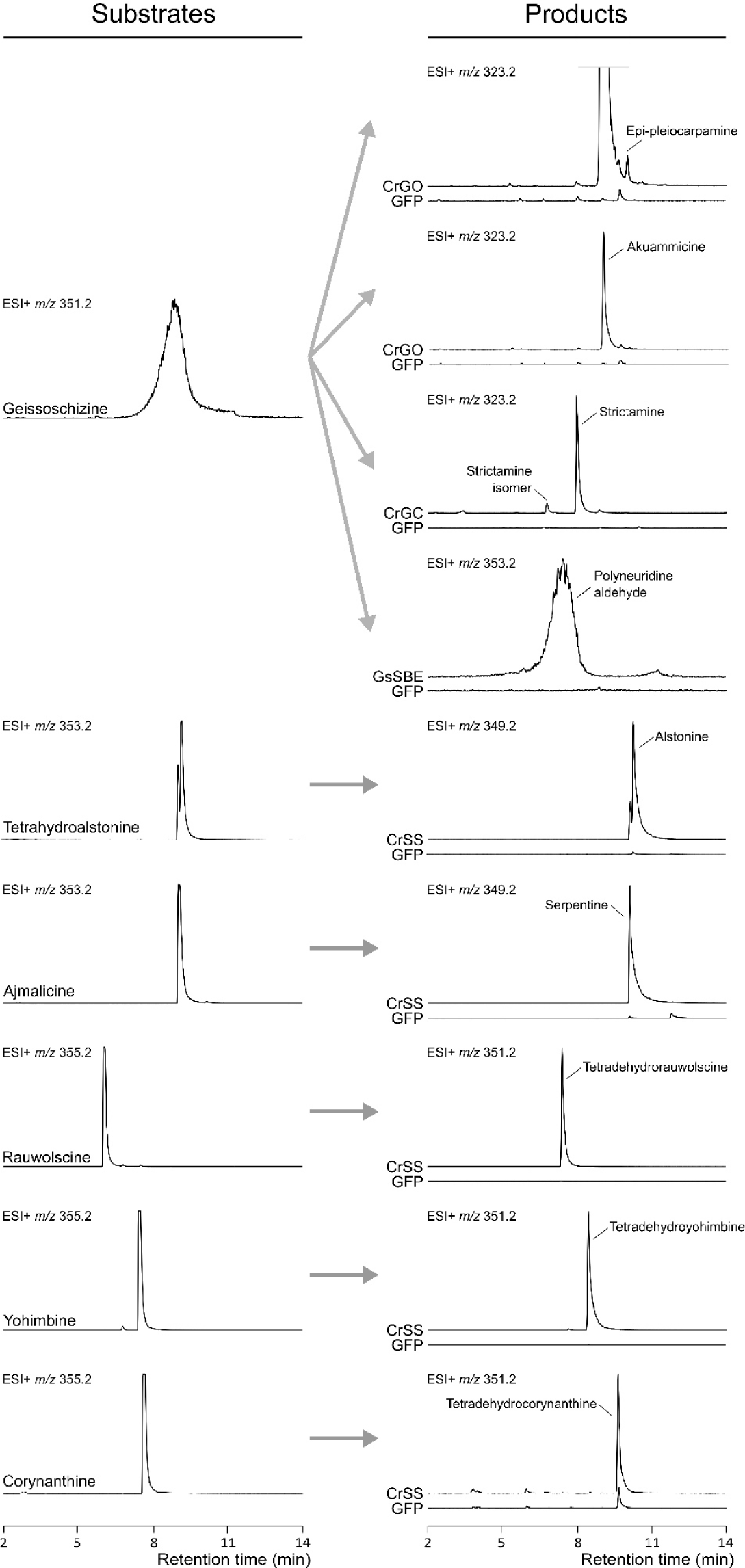
Typical activities of GAS members. CrGC, CrGO, GsSBE and CrSS were used as references for detection of geissoschizine cyclizations (16-*epi*-pleiocarpamine, akuammicine, strictamine isomers and polyneuridine aldehyde) and heteroyohimbane/yohimbane aromatizations (alstonine, serpentine, tetradehydrorauwolscine, tetradehydroyohimbine and tetradehydrocorynanthine). Enzymes were transiently expressed in tobacco leaves and incubated 24h with 55 µM of substrates. LC-MS traces of substrates and products were extracted as ion chromatograms in positive mode at *m/z* 323.2, 349.2, 351.2, 353.2, and 555,2. GFP was used as control.

**Supplemental figure 4:**
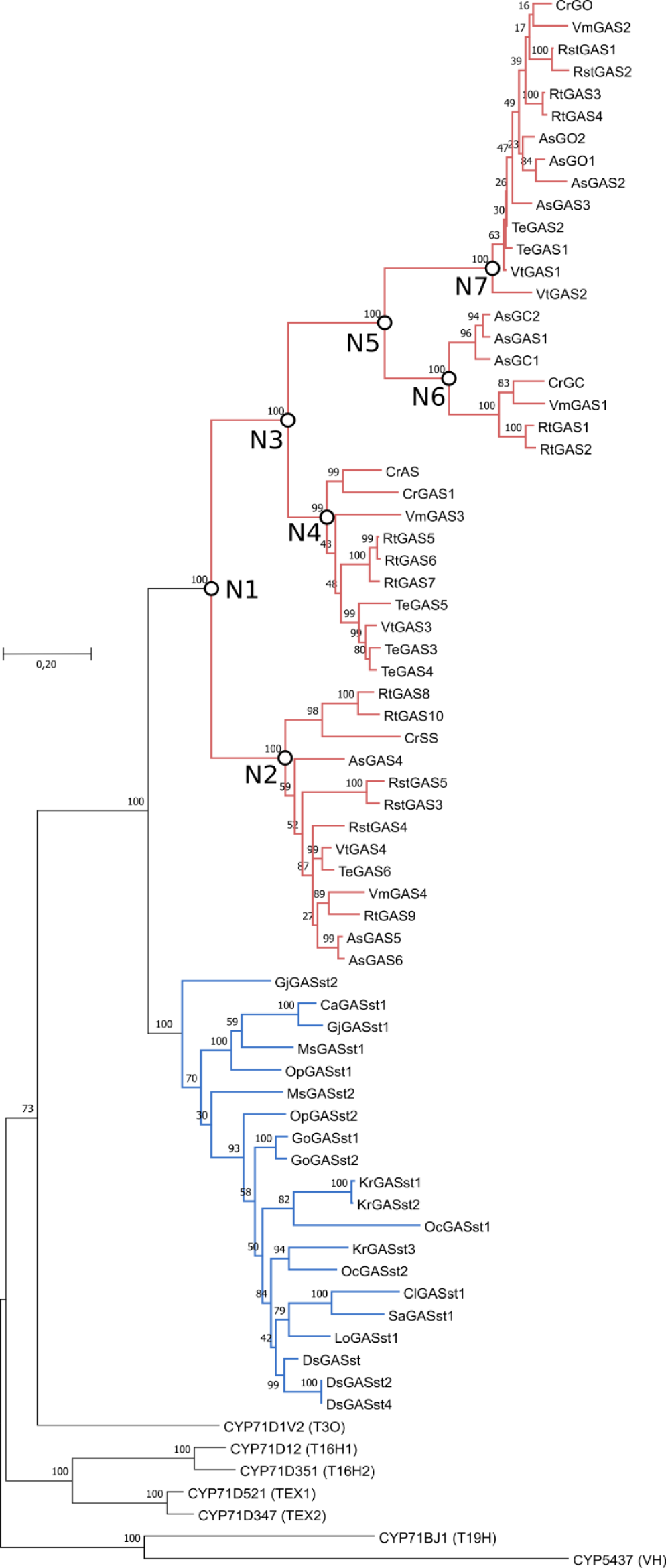
Phylogenetic analysis of GAS and sister GAS predicted from Gentianales genomic resources. Maximum likelihood tree with 1000 bootstrap replicates (%) was inferred using protein sequences curated to remove N-terminal transmembrane domain and indels. Scale bar length represents 0.2 residue substitutions *per* site. GAS clade and sister GAS clade are colored in red and blue, respectively. Nodes targeted for ancestral sequence reconstruction (ASR) are highlighted. N1, LUCA (GAS ancestor); N2, AS/SS group ancestor; N3, SBE/GC/GO ancestor; N4, SBE group ancestor; N5, GC/GO ancestor; N6, GC group ancestor; N7, GO group ancestor.

**Supplemental figure 5:**
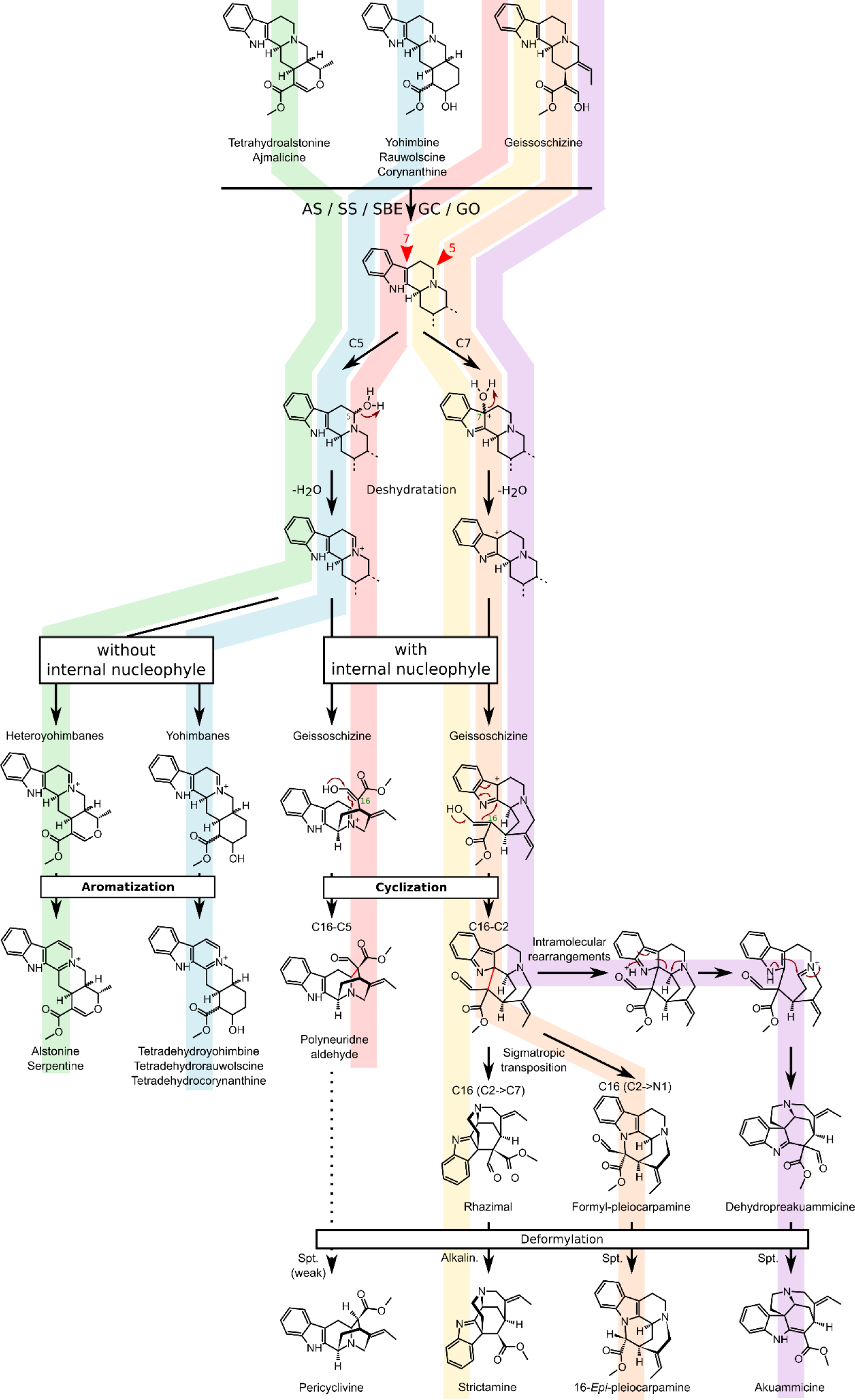
Comparison of proposed GAS catalytic mechanisms. Adapted from Dang et al., 2018, Qu et al., 2018 and Méteignier et al. 2024.

**Supplemental figure 6:**
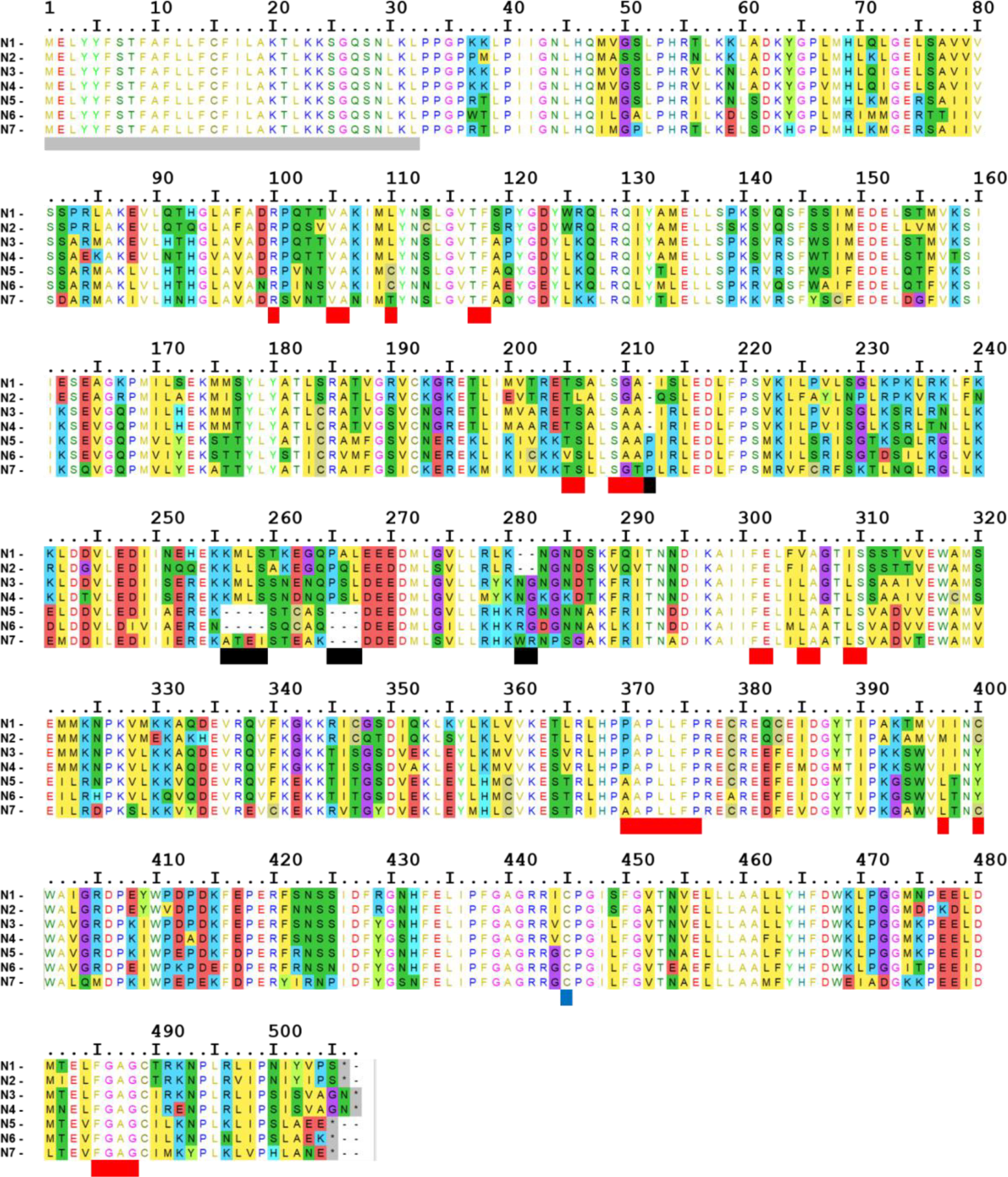
Multiple sequence alignment of ancestral GAS enzymes. The columns of residues without a colored background are strictly conserved between the seven enzymes. The transmembrane domains are shown in gray boxes, indels in black boxes, the cysteine interacting with the heme iron in blue boxes, and the residues of the substrate-binding pocket located above the heme are highlighted in red boxes. N1, LUCA (GAS ancestor); N2, AS/SS group ancestor; N3, SBE/GC/GO ancestor; N4, SBE group ancestor; N5, GC/GO ancestor; N6, GC group ancestor; N7, GO group ancestor.

**Supplemental table 2:**
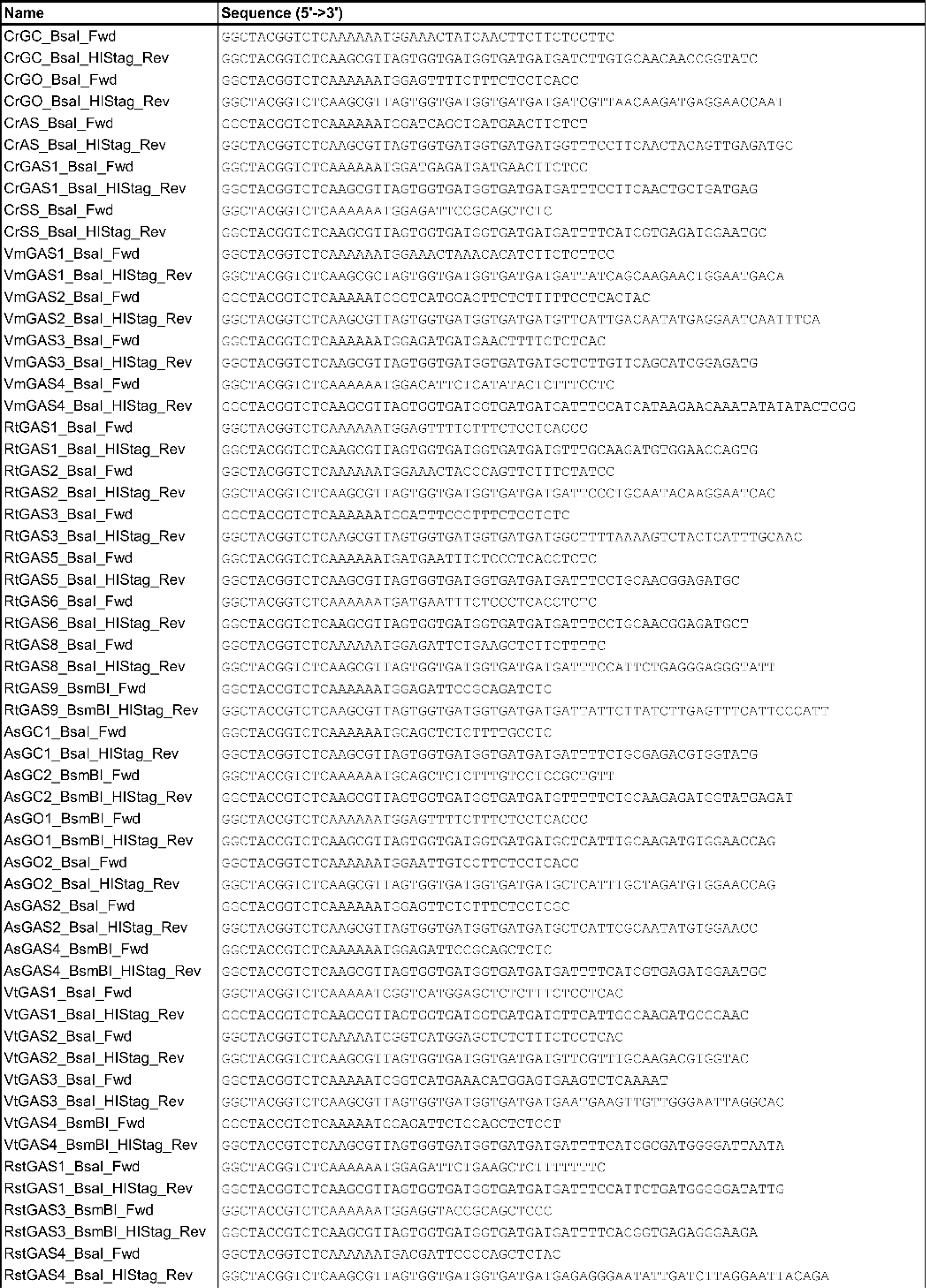
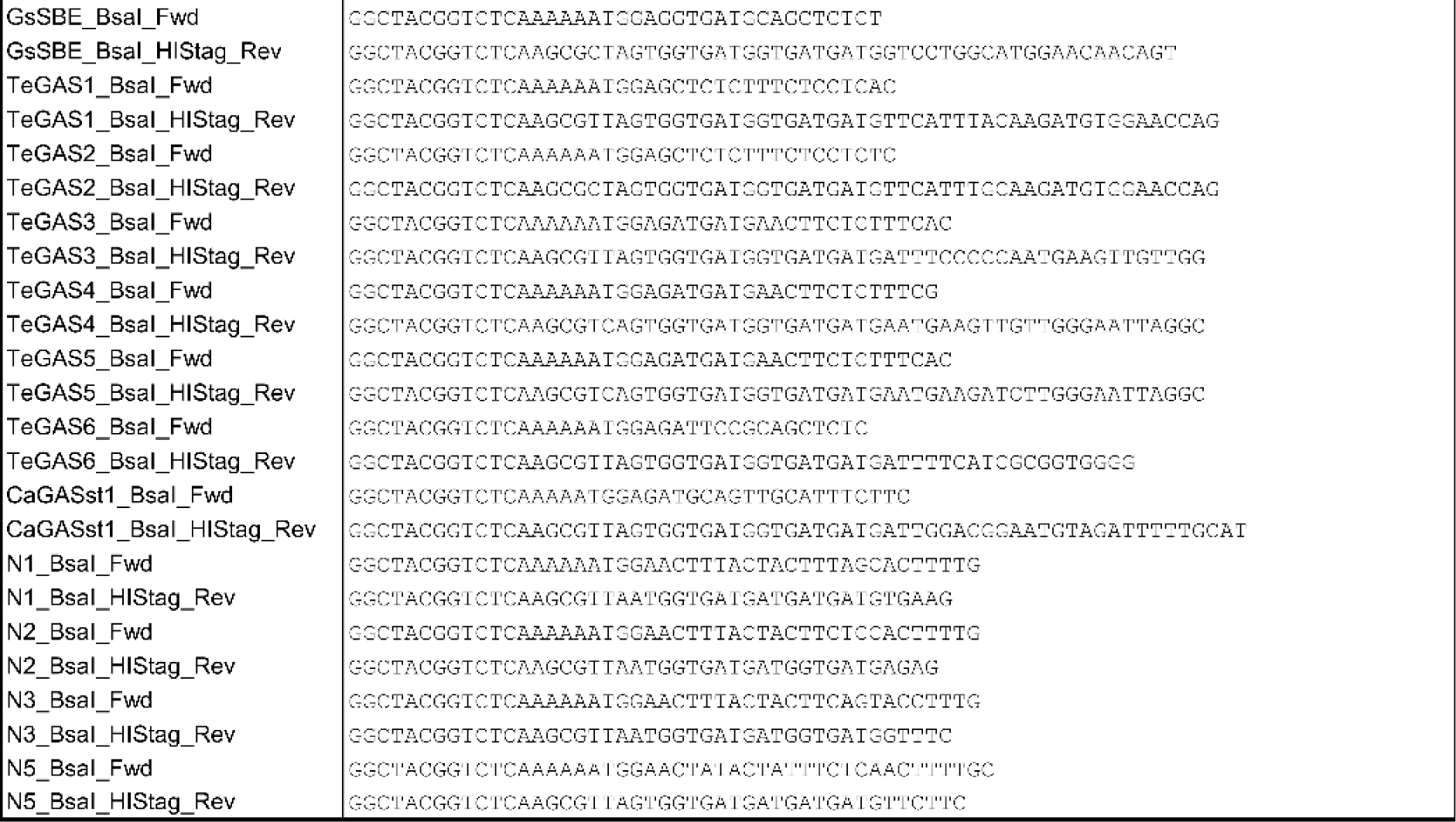
Primers used in this study.

**Supplemental table 3:** Quality parameters of ancestral sequence reconstructed.

| Node | Sequence<br>length | % residues<br>prob > 0,95 | Nb residues<br>prob < 0,6 |
| --- | --- | --- | --- |
| N1 (LUCA) | 470 | 74,1 | 33 |
| N2 | 470 | 89,6 | 6 |
| N3 | 473 | 77,8 | 17 |
| N4 | 473 | 90,7 | 7 |
| N5 | 465 | 71,2 | 32 |
| N6 | 465 | 88,2 | 15 |
| N7 | 469 | 94,9 | 4 |

